# Chemogenetic activation of excitatory neurons alters hippocampal neurotransmission in a dose-dependent manner

**DOI:** 10.1101/595504

**Authors:** Sthitapranjya Pati, Sonali S. Salvi, Mamata Kallianpur, Antara Banerjee, Sudipta Maiti, James P. Clement, Vidita A. Vaidya

**Author notes:** Equal contributing corresponding authors, **Address correspondence to:** Dr. Sthitapranjya Pati, Department of Biological Sciences, Tata Institute of Fundamental Research, 1 Homi Bhabha Road, Mumbai 400005, India, Dr. James P. Clement, Neuroscience Unit, Jawaharlal Nehru Centre for Advanced Scientific Research, Jakkur, Bangalore 560 064, India., Dr. Vidita A. Vaidya, Department of Biological Sciences, Tata Institute of Fundamental Research, 1 Homi Bhabha Road, Mumbai 400005, India.

## Abstract

Designer Receptors Exclusively Activated by Designer Drugs (DREADD)-based chemogenetic tools are extensively used to manipulate neuronal activity in a cell-type specific manner. Whole-cell patch-clamp recordings indicate membrane depolarization, coupled with increased neuronal firing rate, following administration of the DREADD ligand, Clozapine-N-Oxide (CNO) to activate the Gq-coupled DREADD, hM3Dq. Although hM3Dq has been used to enhance neuronal firing in order to manipulate diverse behaviors, often within thirty minutes to an hour post-CNO administration, the physiological effects on excitatory neurotransmission remain poorly understood. We investigated the influence of CNO-mediated hM3Dq DREADD activation on distinct aspects of hippocampal excitatory neurotransmission at the Schaffer collateral-CA1 synapse in hippocampal slices derived from mice expressing hM3Dq in Ca^2+^/calmodulin dependent protein kinase α (CamKIIα)-positive excitatory neurons. Our results indicate a clear dose-dependent effect on fEPSP slope, with no change noted at the lower dose of CNO (1 µM) and a significant, long-term decline in fEPSP slope observed at higher doses (5-20 µM). Further, we noted a robust theta burst stimulus (TBS) induced long-term potentiation (LTP) in the presence of the lower CNO (1 µM) dose, which was significantly attenuated at the higher CNO (20 µM) dose. Whole-cell patch clamp recording revealed both complex dose-dependent regulation of excitability, and spontaneous and evoked activity of CA1 pyramidal neurons in response to hM3Dq activation across CNO concentrations. Our data indicate that CNO-mediated activation of the hM3Dq DREADD results in dose-dependent regulation of excitatory hippocampal neurotransmission, and highlight the importance of careful interpretation of behavioral experiments involving chemogenetic manipulation.

## Introduction

Spatio-temporal control and manipulation of neuronal activity is an essential tool to understand the causal relationship between behavior and underlying neuronal circuits. Presently, two major technologies are used to perturb neuronal function; (1) optogenetic tools which are light-activated ion channels (Britt and Bonci, 2013; Kim et al., 2017; Xie et al., 2013) and (2) chemogenetic tools which are modified G protein-coupled receptors (GPCRs) that can be activated by a pharmacologically inert ligand Clozapine-N-oxide (CNO) (Nichols and Roth, 2009; Pei et al., 2008; Roth, 2016; Urban and Roth, 2015). While optogenetics is immensely useful for excitation/ inhibition of neuronal activity in behavioral time scales ranging from milliseconds to several minutes, chemogenetic activation/ inhibition of neuronal circuits is predominantly the method of choice for more long-term perturbations ranging from a few minutes to chronic manipulation across days (Kim et al., 2017; Roth, 2016). While both optogenetic and chemogenetic tools can be combined with inducible genetics to manipulate neuronal activity in a circuit/ cell type-specific manner in precise temporal windows, optogenetics is disadvantageous for chronic manipulation due to problems such as overheating of tissue and the need to implant optical fibers by invasive surgery (Britt and Bonci, 2013; Kim et al., 2017; Roth, 2016).

The Designer Receptors Exclusively Activated by Designer Drugs (DREADD) hM3Dq, which is an engineered human muscarinic receptor, is routinely used to activate neurons (Alexander et al., 2009; Roth, 2016). The hM3Dq DREADD is coupled to Gq-signaling and like other Gq-coupled receptors, application of its agonist CNO increases accumulation of inositol monophosphate (IP1) and mobilizes intracellular calcium (Alexander et al., 2009; Armbruster et al., 2007; Guettier et al., 2009; Roth, 2016). CNO administration leads to depolarization of membrane potential along with increased firing rate in hippocampal CA1 pyramidal neurons (Alexander et al., 2009), raphe serotonergic neurons (Urban et al., 2016), and arcuate nucleus AgRP neurons (Krashes et al., 2011). Although hM3Dq has been used as a tool to increase neuronal activity in several circuits including those responsible for feeding (Atasoy et al., 2012; Krashes et al., 2011), locomotion (Kozorovitskiy et al., 2012), energy expenditure (Kong et al., 2012), memory (Garner et al., 2012), and social behaviors (Penagarikano et al., 2015), detailed neurophysiological consequences of its activation still remain poorly characterized.

Many other Gq-signaling-coupled metabotropic receptors like the metabotropic glutamate receptor 1 and 5 (mGluR1/5), muscarinic acetylcholine receptor 1 and 5 (M1/5) are known to modulate neuronal cells and circuits in distinct fashion at different doses of ligand (Caruana et al., 2011; Gladding et al., 2009; Kumar, 2010; Volk et al., 2007). Although a wide range of dosage spanning from 0.5 to 200 µM of CNO has been used to record hM3Dq mediated electrophysiological effects in acute slices (Hurni et al., 2017; Mahler et al., 2014), possible dose-dependent variation in neuronal activity has not been studied. Further, most behavioral assays are carried out from thirty minutes to one-hour post-CNO administration (Roth, 2016), a timescale in which the physiological effects of CNO remain poorly understood.

In the current study, we have investigated the effects of acute CNO mediated activation of the hM3Dq receptor in the Ca^2+^/calmodulin dependent protein kinase α (CamKIIα)-positive excitatory neurons on hippocampal neurotransmission pre and thirty minutes post-CNO administration. Using the Schaffer collateral pathway as the model circuit, we show that acute hM3Dq activation in excitatory neurons produces distinctly different dose-dependent effects on field currents, paired-pulse facilitation, neuronal excitability, spontaneous postsynaptic currents (sPSC), and ionotropic glutamate receptor-mediated currents. We also observe a dose-dependent increase in intracellular calcium levels in cultured hippocampal neurons following CNO administration. Taken together, our results suggest that hM3Dq activation in hippocampal CamKIIα-positive excitatory neurons produces clear dose-dependent effects on neurotransmission, highlighting the importance for careful interpretation of behavioral studies using chemogenetic activation.

## Materials and Methods

### Animals

CamKIIα-tTA transgenic mice were received as a gift from Dr. Christopher Pittenger, Department of Psychiatry (Mayford et al., 1996), Yale School of Medicine. TetO-hM3Dq mice (Cat. No. 014093; Tg(tetO-CHRM3*)1Blr/J) were purchased from Jackson Laboratories, USA. CamKIIα-tTA and TetO-hM3Dq bigenic mice were bred, and genotypes of double positive animals were confirmed using PCR based analysis. The background strain C57Bl6/J was used for control experiments. Male mice were used for all experiments. All animals were maintained in a 12-hour light-dark cycle and provided with *ad libitum* access to food and water. All experimental procedures were carried out following the Committee for the Purpose of Control and Supervision of Experiments on Animals (CPCSEA), Government of India and were approved by both the TIFR and JNCASR institutional animal ethics committee.

### Drug administration

Clozapine-N-oxide (CNO; Tocris, UK) was dissolved in artificial cerebrospinal fluid (aCSF). This solution was continuously bubbled with 95% O_2_ and 5% CO_2_ and was circulated through the slice chamber during drug administration.

### Immunofluorescence and confocal imaging

As the TetO-hM3Dq mouse has a hemagglutinin (HA) tag, we performed an immunofluorescence staining to visualize the hM3Dq expression in the hippocampus and cortex. The CamKIIα-tTA::TetO-hM3Dq double positive or genotype-control animals (single positive for either CamKIIα-tTA or TetO-hM3Dq) were sacrificed by transcardial perfusion with saline (0.9% NaCl) followed by 4% paraformaldehyde. Coronal brain sections (40 µm) were generated using a vibrating microtome (Leica, Germany) and subjected to immunofluorescence. Following permeabilization at room temperature in phosphate buffered saline with 0.4% Triton X-100 (PBSTx) for one hour, the sections were subjected to blocking solution (1% Bovine Serum Albumin (Roche, 9048-49-1), 5% Normal Goat Serum (Thermoscientific, PI-31873) in 0.4% PBSTx) at room temperature for one hour. Sections were then incubated in the primary antibody, rabbit anti HA (1:250; Rockland, 600-401-384, USA) for 4 days at 4°C which was followed by three washes with 0.4% PBSTx of fifteen minutes each at room temperature. This was followed by incubation with the secondary antibody, goat anti-rabbit IgG conjugated to Alexa Fluor 568 (1:500; Invitrogen, A-11079, USA) for two and a half hours at room temperature. The sections were mounted onto slides with Vectashield Antifade Mounting Medium with DAPI (Vector, H-1200, USA). The sections were imaged using an LSM5 exciter confocal microscope (Zeiss, Germany) using identical acquisition settings for the sections from double positive and genotype-control animals.

### Primary hippocampal neuron culture

Hippocampi were extracted from pups of CamKIIα-tTA::TetO-hM3Dq at postnatal day 0 and tail-clips were collected for genotyping. Hippocampi were dissected in HBSS-HEPES (300mM) buffer and incubated with 0.1% trypsin-EDTA (Invitrogen, USA) for 10 min. Neurons were dissociated in Neurobasal medium supplemented with 2% B27 supplement and 0.5mM L-glutamine (Invitrogen, USA). Cells were plated on poly-D-Lysine (Sigma) coated plates at a density of 10^6^ cells/well in neurobasal medium. Calcium imaging experiments were carried out at day in vitro (DIV) 6-8.

### Preparation of hippocampal slices

CamKIIα-tTA::TetO-hM3Dq or C57Bl/6J mice were sacrificed by cervical dislocation in accordance with the guidelines of the JNCASR animal ethics committee. Following decapitation, the brain was immediately transferred to ice cold sucrose cutting solution (189 mM Sucrose, 10 mM D-glucose, 26 mM NaHCO_3_, 3 mM KCl, 10 mM MgSO_4_.7H_2_O, 1.25 mM NaH2PO_4_, and 0.1 mM CaCl_2_) which was bubbled continuously with 95% O_2_ and 5% CO_2_. Following dissection and removal of the cerebellum in a petri dish containing an ice-cold cutting solution, the brain was glued onto a brain holder which was placed in a buffer tray containing ice-cold cutting solution. Subsequently, 300 µm horizontal sections were obtained using a vibrating microtome (Leica, VT-1200, Germany). The sections were then transferred to a petri dish containing aCSF (124 mM NaCl, 3 mM KCl, 1 mM MgSO_4_.7H_2_O, 1.25 mM NaH_2_PO_4_, 10 mM D-glucose, 24 mM NaHCO_3_, and 2 mM CaCl_2_) at room temperature following which the hippocampus and the overlying cortex was gently dissected. Slices were transferred to a chamber on a nylon mesh containing aCSF bubbled with 95% O_2_ and 5% CO_2_ at 37°C. It was incubated for thirty minutes to one hour to ensure stable electrophysiological responses. The slices could be maintained in a healthy state for up to eight hours and were transferred to the recording chamber as required.

### Extracellular and intracellular recording rig

The aCSF was pre-heated to 34°C using an online Peltier controlled temperature control system (ThermoClamp-1, Automate Scientific, USA) and circulated through the slice recording chamber (Scientifica, UK) at 1-2 mL min^−1^ using a combination of peristaltic pump (BT-3001F, longer precision pump Co. Ltd., China) and gravity feed. An Ag/AgCl reference wire and a thermocouple to provide feedback to the temperature control system were submerged in the recording chamber containing aCSF. All stimulating and recording electrodes were placed in the slice approximately 45° to the vertical and were controlled using a micromanipulator (# 1U RACK; Scientifica, UK) which allowed movement in all three axes for correct positioning on the slice. The slice and the electrodes were visualized using an upright microscope (Slicescope pro 6000 Scientifica, UK) with specialized optics to visualize deep tissue. The recording set up was mounted on an anti-vibration table (#63P-541; TMC, USA) and were enclosed in a Faraday cage. The electrical noise was eliminated by grounding all electrical connections to a single ground point in the amplifier.

### Recording techniques

#### Recording electrodes

Recording electrodes were pulled from borosilicate glass capillaries (# 30-0044/ GC120F-10; Harvard apparatus, UK) using a horizontal micropipette puller (# P97, Sutter Instruments Co., USA). While intracellular patch electrodes (5-7 MΩ) were filled with potassium gluconate (KGlu) internal solution (130 mM KGlu, 20 mM KCl, 10 mM HEPES free acid, 0.2 mM EGTA, 0.3 mM GTP-Na salt, 4 mM ATP-Mg salt; osmolarity adjusted to 280-310 mOsm), electrodes used for extracellular field recording (3-5 MΩ) were filled with aCSF. The microelectrodes were mounted on the electrode holder (Scientifica, UK) so that the Ag/AgCl recording wire was in touch with the pipette solution. This holder was mounted to on a headstage which was connected to the amplifier. All signals were amplified using a Multiclamp-700B (Molecular Devices, USA).

#### Extracellular field recording

The measurement of field excitatory postsynaptic potential (fEPSP) was carried out from the CA1 *stratum radiatum* and was measured as the potential difference between the recording electrode and the bath electrode.

#### Whole cell patch-clamp recording

Whole cell patch-clamp recording was carried out from somata of CA1 pyramidal neurons. A positive pressure was applied to the patch pipette filled with KGlu intracellular recording solution using a tube attached to the pipette holder. With the current and voltage offset to zero, the resistance of the electrode was checked by applying a test pulse and measuring current deflection according to Ohm’s law. The positive pressure was released when the pipette tip touches the cell surface which was confirmed both by visualization under the microscope and a change in resistance. This was followed by application of a negative pressure through gentle suction to form a tight seal, indicated by a large increase in resistance (> 1 GΩ). The slow and fast capacitance were adjusted following which the patch of the membrane was ruptured by application of a gentle negative pressure and if required a strong voltage pulse. The cells with membrane potential less than −55 mV and a series resistance in the range of 5-25 MΩ were considered for future experiments.

#### Stimulation protocol

The Schaffer collateral-commissural fiber pathway was stimulated using a concentric bipolar stimulating electrode (Outer diameter: 125 µm, Inner diameter: 25 µm Platinum/Iridium, CBARC75, FHC, USA). For extracellular recordings, stimulus of a square-wave pulse of 20-100 µs in duration and 20-200 µA in amplitude was applied using an isolated direct current stimulation box (Digimeter, UK). For all experiments, a paired-pulse stimulus was delivered at 50 ms interval per sweep with 20s inter-sweep intervals and a potentiated paired-pulse was used to confirm proper placement of electrodes in Schaffer collateral. Following the acquisition of input-output (I-O) characteristic and paired-pulse ratio (PPR), a stable baseline of the fEPSP slope was established. While I-O curves were acquired at a fixed 20-40 µs duration and increasing stimulus intensity from 0-300 µA, the PPR was measured at a 10s inter sweep interval with inter-stimulus intervals ranging from 10-1000 ms in a quarter-logarithmic scale. CNO (1, 5, 10, or 20 µM) containing aCSF was administered and a constant time course was acquired for 60 min. We selected the lowest (1 µM) and highest (20 µM) CNO dosage for further experiments. I-O response and PPR were measured before and thirty minutes following CNO administration while a constant time course of stimulus-induced fEPSP was recorded. In a separate set of slices, theta burst stimulation (TBS) was used to induce long-term potentiation (LTP) following either a low-dose (1 µM) or a high dose (20 µM) of CNO. For the TBS-induced LTP-protocol, thirteen blocks of a four-pulse 100 Hz stimuli were applied with an inter-sweep interval of 200 ms (5 Hz).

All measurements were done before and thirty minutes following CNO (1 µM or 20 µM) administration, except for AMPAR and NMDAR-mediated currents for which a time-course was acquired across thirty minutes. Intrinsic cellular properties such as resting membrane potential, input resistance, tau, sag voltage, were calculated by a 500 ms, −100 pA to 180 pA, 7-step hyperpolarizing or depolarizing current injection with an inter sweep interval of 10s. To confirm cell identity by observing the shape of the action potential and calculate the action potential threshold, a current of up to 2 nA for 2 ms was injected. Spontaneous currents were measured by holding the cell in voltage-clamp mode at −70 mV. To measure evoked response, the Schaffer collateral-commissural pathway was stimulated using a concentric bipolar stimulating electrode (Outer diameter: 125 µm, Inner diameter: 25 µm Platinum/Iridium, CBARC75, FHC, USA) and cells were voltage clamped either at −70 mV for AMPAR-mediated currents or +40 mV for NMDAR-mediated currents.

#### Calcium imaging

In order to detect intracellular calcium levels following administration of CNO, we used the UV-excitable calcium-sensitive ratiometric dye Indo-1 AM, Cell-permeable (I1226, Thermo Fischer Scientific, USA). A custom-built two-photon set up with a Ti:Sapphire laser (MaiTai DeepSee, Spectra Physics, USA) coupled to a confocal microscope (LSM 710, Carl Zeiss, Jena, Germany), as described earlier (Das et al., 2017) was used for imaging (excitation: 730-735 nm). A microscope incubator stage (Okolab, Italy) was used to maintain 37°C and 5% CO_2_. A combination of liquid copper sulfate filter (for infrared) and 400/30 nm bandpass filter was used to detect the emission of the calcium-bound dye. Primary hippocampal neurons (DIV 6-8) were incubated in 10 µM Indo-1 AM made in 1X Hank’s balanced salt solution (HBSS, 14175-079, Gibco, Life Technologies, USA) supplemented with 100 nM CaCl_2_ for twenty minutes at 37°C, 5% CO_2_. Following three brief washes with 1X HBSS, a baseline fluorescence of ten minutes was acquired using a 40X objective. Following this, CNO (1 µM or 20 µM) was added and the neurons were tracked for thirty minutes, with image stacks acquired every two minutes.

### Data analysis

#### Extracellular field recording data

All extracellular field recording data were analyzed off-line using Clampfit 10.5 (Molecular Devices, USA). The slope of the rising phase of the fEPSP was used as a measure of the size of fEPSP to eliminate the possibility of contamination with population spikes. For the I-O measurements, the fEPSP slope was calculated with increasing stimulus intensity (0-300 µA). Following a paired-pulse stimulation of the Schaffer collateral pathway, a potentiation in fEPSP slope was observed. The paired pulse ratio was calculated as the ratio of the fEPSP slopes following the first stimulus to the paired stimulus following inter-pulse intervals of 10-1000 ms in a quarter-logarithmic scale.

#### Whole cell voltage/ current clamp data

All whole cell voltage/ current clamp data were analyzed off-line using Clampfit 10.5 (Molecular Devices, USA). The intrinsic membrane properties were measured using voltage deflection traces from 100 pA hyperpolarizing current pulse. The input resistance was calculated by applying Ohm’s law, R = V/I. Where V = stable state voltage and I = injected current (100 pA). The membrane time constant (τ_m_) was obtained by fitting the voltage decay with a single exponential Ae^-t/τ^ and calculating the decay constant. The sag was calculated by subtracting the steady state voltage deflection from the peak negative-going voltage. I-O curve was obtained for the number of spikes fired in response to increasing amplitude of current injection (0-180 pA). Following this, the accommodation index was calculated as the ratio of the maximum inter-spike interval (including the interval from the last spike to the end of the current injection) to the first inter-spike interval. The action potential threshold was defined as the voltage where the rate of change of voltage with respect to time (dV/dt) exceeded 10 Vs^-1^. The spontaneous current amplitude and inter-event intervals were calculated using an automated event detection algorithm using Mini Analysis program (Synaptosoft Inc., USA). An average of at least 200 events was detected from each 5 min trace pre and thirty minutes post-CNO administration and then subjected to statistical analysis.

AMPAR-mediated current was calculated as the peak amplitude of evoked current when the cell was voltage clamped at −70 mV and NMDAR-mediated currents were obtained as the average current 80-100 ms following the time of peak response when the cell was voltage clamped at +40 mV. The NMDAR-mediated current decay kinetics were calculated by fitting a double exponential function in decay phase of the NMDAR-mediated current and applying the results to the following equation: τ_w_ = [I_f_ /(I_f_ + I_s_)] x τ_f_ + {I_s_ / (I_f_ + I_s_)} x τ_s_. Where, τ_w_ is the weighted tau, I_f_ and I_s_ are the amplitudes of fast and slow currents, τ_f_ and τ_s_ are decay constants of fast and slow currents respectively (Rumbaugh and Vicini, 1999).

#### Calcium imaging data

Calcium imaging data were analyzed using ImageJ. Following a maximum intensity projection, the time series images were corrected for any horizontal drift using scale-invariant feature transfer (SIFT) algorithm (Lowe, 2004). A region of interest was drawn around the neurons of interest and average intensity values were calculated. Fluorescent intensities were normalized to the baseline following which statistical analysis was performed.

#### Statistics

Linear regression was used to analyze the time course data, while to analyze time-binned datasets, one-way analysis of variance (ANOVA) was performed followed by Bonferroni post-hoc analysis (GraphPad Prism, Graphpad Software Inc., USA). Data are expressed as mean ± standard error of mean (S.E.M) and statistical significance was determined at *p* value < 0.05. The spontaneous current data was analyzed using MATLAB (Mathworks, USA). The amplitude and inter-event interval were converted to corresponding cumulative probability distributions, and then subjected to Kolmogorov-Smirnov two-sample comparison. Statistical significance was set at *p* value < 0.001.

## Results

### Selective expression of hM3Dq DREADD in forebrain excitatory neurons

The selective expression of hM3Dq DREADD in Ca^2+^/calmodulin-dependent protein kinase α (CamKIIα)-positive excitatory neurons in the forebrain was achieved by generating a bigenic mouse created by crossing CamKIIα-tTA and TetO-hM3Dq mouse lines (Fig. 1A-B). We performed immunofluorescence analysis to detect the HA-tag and thus visualize the expression of hM3Dq in hippocampal subdivisions, including the CA1 (Fig. 1C) and dentate gyrus (DG; Fig. 1D) subfields, and also in the neocortex (Fig. 1E) of CamKIIα-tTA::TetO-hM3Dq animals. Analysis in single positive mice CamKIIα-tTA or TetO-hM3Dq mice indicated no HA-Tag immunofluorescence in the CA1, DG hippocampal subfields or the neocortex (Fig. 1F-H).

**Fig. 1.**
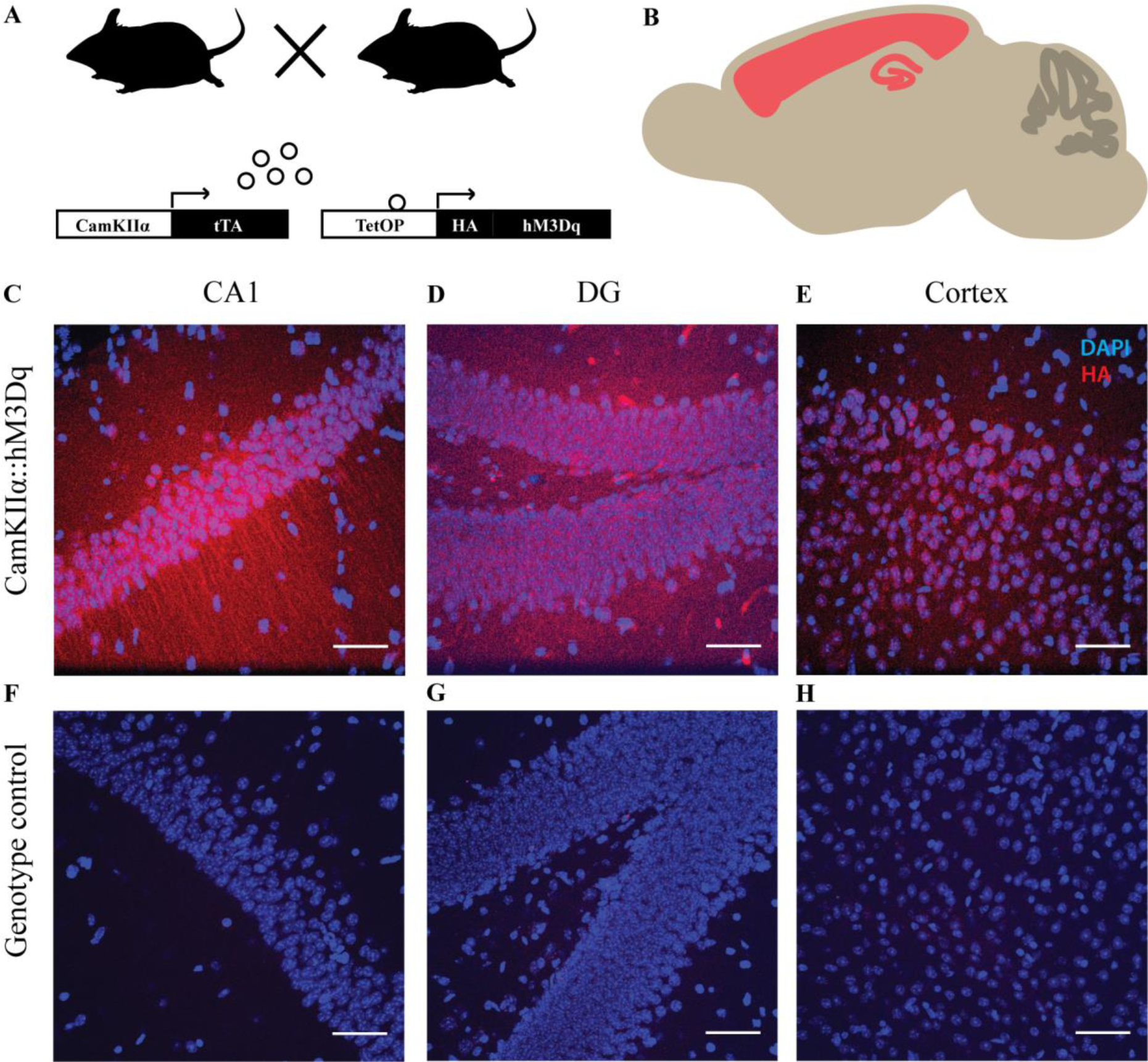
Selective expression of hM3Dq DREADD in forebrain excitatory neurons. (A) Shown is a schematic of experimental strategy. hM3Dq DREADD was selectively expressed in the CamKIIα-positive excitatory neurons in the forebrain. (B) Shown is a schematic of the extent of expression. Representative confocal images showing expression of hM3Dq identified by fluorescent immune-staining of HA-tag in the CA1 (C), DG (D), and cortex (E). No HA-tag immunoreactivity was observed in the genotype-control animals (F-H). Blue: DAPI, Red: HA-Tag. Scale bar = 50 µm.

### Acute chemogenetic activation of hippocampal excitatory neurons differentially regulates field transmission in Schaffer collaterals in a dose-dependent manner

To investigate effects of chemogenetic activation of CamKIIα-positive excitatory neurons on hippocampal neurotransmission, we performed time course measurements of field excitatory post synaptic potential (fEPSP) slope in the stratum radiatum in response to stimulation of Schaffer collateral-commissural fiber pathway (Fig. 2A). After establishing a stable baseline, we perfused aCSF containing different dosages of CNO following which fEPSP was recorded for sixty minutes. We did not find any significant change in the fEPSP slope over time at the lowest dose of CNO (1 µM; Sup Fig. 1A, n = 3). Interestingly, we observed a decline in fEPSP slope over time with bath application of 5 µM (Sup Fig. 1B, n = 3), 10 µM (Sup Fig. 1C, n = 4), and 20 µM (Sup Fig. 1D, n = 4) CNO. To rule out potential non-specific effects mediated by CNO, we performed the same experiment using acute hippocampal slices from the background strain, C57Bl/ 6J. We did not note any change in fEPSP slope over time with bath application of either 1 µM (Sup Fig. 1E, n = 3) or 20 µM (Sup Fig. 1F, n = 3) CNO. For further investigation of dose-dependent effects of CNO mediated activation of CamKIIα-positive excitatory neurons in the hippocampus, we chose the lowest (1 µM) and the highest (20 µM) dose of CNO. As evident from these results, the CNO mediated decline in fEPSP slope reached steady state at around thirty minutes post-CNO administration. Therefore, we performed time course measurements for thirty minutes in a separate cohort of animals. We observed no change in fEPSP time course with the low dose (1 µM) of CNO and a significant decline in fEPSP slope over time following high dose (20 µM) of CNO administration (Fig. 2B). The fEPSP slope time course was significantly lower at 20 uM CNO (Linear regression followed by ANCOVA, *p* < 0.0001, n = 5 for 1 µM and n = 6 for 20 µM) as compared to the 1 µM CNO treated slices. We next measured the input-output response of the pathway before CNO treatment (Pre-CNO) and 30 minutes post-CNO treatment. At low dose, we did not observe any significant difference in fiber volley (FV) peak (Fig. 2C, n = 5) and fEPSP slope (Fig. 2D, n = 5) in response to increasing stimulus intensity. At the high dose, we did not find any significant difference in the FV peak in response to increasing stimulus intensity (Fig. 2E, n = 5). Interestingly, we found that the fEPSP slope in response to increasing stimulus intensity was lower after thirty minutes of 20 µM CNO administration (Fig. 2F, linear regression followed by ANCOVA, *p* = 0.0003, n = 5). For the input-output response curve, we plotted fEPSP slope against FV peak at corresponding stimulus intensity where we did not find any change with thirty minutes of low dose of CNO administration (Sup Fig. 2A, n = 5). The fEPSP slope in response to corresponding FV peak was reduced with thirty minutes administration of high dose of CNO (Sup Fig. 2B, n = 5). Paired-pulse facilitation in the Schaffer collaterals has been shown to be mediated by pre-synaptic mechanisms. To assess possible pre-synaptic effects of CNO administration, we measured paired-pulse facilitation before and thirty minutes following CNO administration. Strikingly, we found a potentiated paired-pulse facilitation at thirty minutes following 1 µM CNO administration (Fig. 2G, linear regression followed by ANCOVA, *p* = 0.041, n = 9). We did not find any significant change in paired-pulse ratio with increasing interpulse interval after thirty minutes of 20 µM CNO administration (Fig. 2H, n = 10) as compared to pre-CNO measurements.

**Fig. 2.**
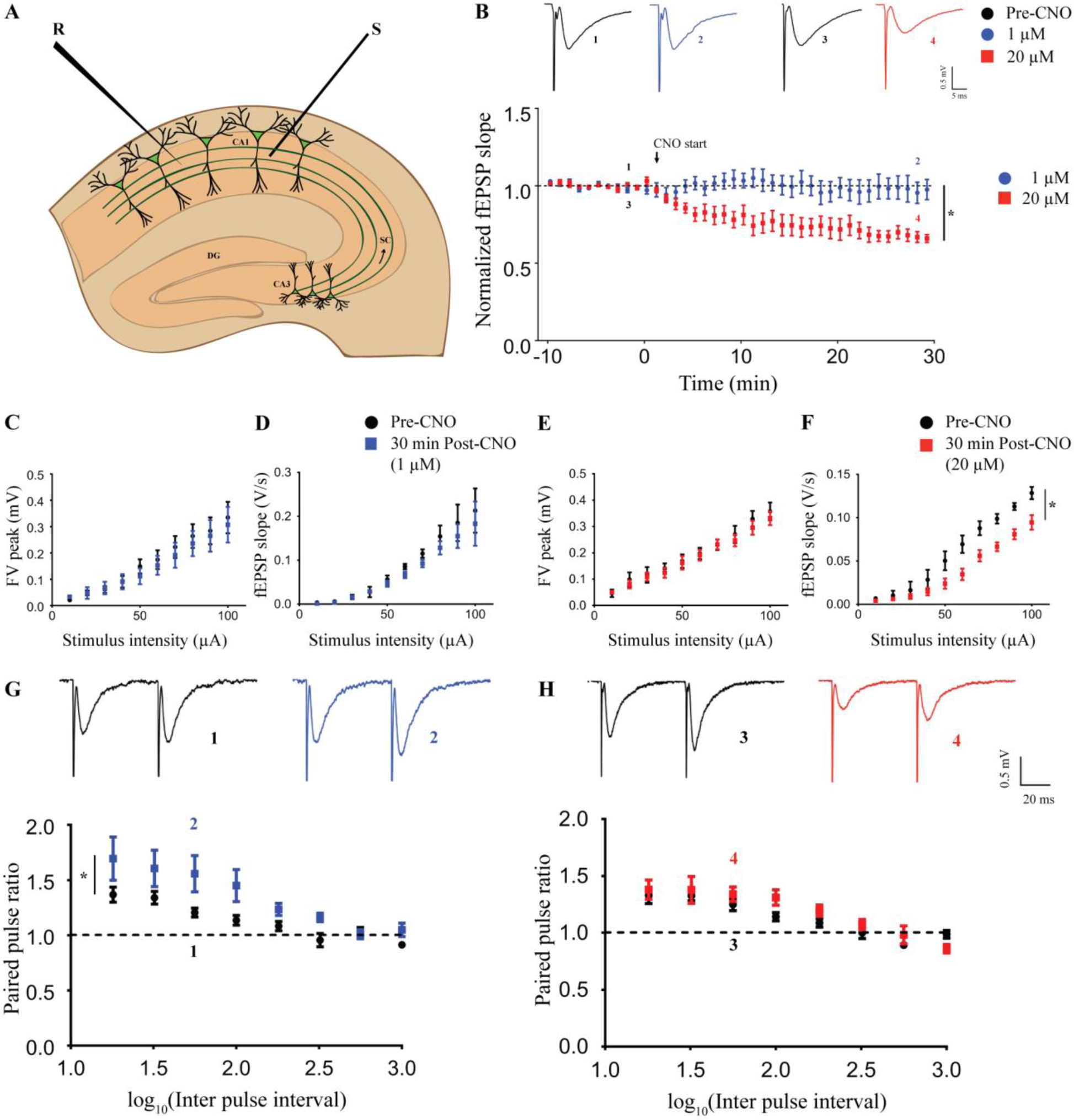
Acute chemogenetic activation of hippocampal excitatory neurons differentially regulates field transmission in Schaffer collaterals in a dose-dependent manner. (A) Shown is a schematic of placement of stimulating and recording electrode in the Schaffer collaterals. (B) Shown are representative fEPSP traces before (1) and thirty minutes (2) following 1 µM CNO treatment and representative fEPSP traces before (3) and thirty minutes (4) following 20 µM CNO treatment. Bath application of CNO did not affect fEPSP time course at 1 µM, whereas it led to long-term depression of fEPSP at 20 µM. Stimulus intensity: 50-80 µA; 20-30 µs. The FV peak was not altered with increasing stimulus intensity either at 1 µM (C) or 20 µM (E). The fEPSP slope was not different at 1 µM (D), but significantly lower in response to increasing stimulus intensity at 20 µM (F). (G) Shown are representative paired-pulse evoked fEPSP traces before (1) and thirty minutes (2) following 1 µM CNO treatment. A significant potentiation of paired-pulse ratio was observed (G) following thirty minutes of 1 µM CNO bath application. (H) Shown are representative paired-pulse evoked fEPSP traces before (3) and thirty minutes (4) following 20 µM CNO treatment. No significant alteration in the paired-pulse ratio was noted following thirty minutes of 20 µM CNO bath application (H). Results are expressed as the mean ± S.E.M. \**p* < 0.05 as compared between 1 µM and 20 µM CNO treatment (linear regression followed by ANCOVA). S – stimulating electrode, R – recording electrode.

The above results indicate that acute chemogenetic activation of CamKIIα-positive excitatory neurons produces a long-term decline of fEPSP in the Schaffer collateral at a high dose (20 µM) of CNO, while no significant effect was observed on the fEPSP time course at the low dose (1 µM). Further, our results from the paired-pulse facilitation experiments indicate a putative potentiation of pre-synaptic responses in the Schaffer collaterals following chemogenetic activation of CamKIIα-positive excitatory neurons using a low dose (1 µM) of CNO.

### Dose-dependent effects of acute chemogenetic hM3Dq activation on theta burst stimulation induced LTP in hippocampal slices

To assess whether hippocampal plasticity is modulated by hM3Dq DREADD activation at the low and high dose of CNO, we induced long-term potentiation (LTP) following thirty minutes of either 1 µM or 20 µM CNO. We used a weak, physiologically relevant theta burst stimulation protocol (TBS). Interestingly, we observed a robust long-term potentiation of fEPSP slope in the slices pre-treated with 1 µM CNO for thirty minutes (Fig. 3A-B). However, in hippocampal slices pre-treated with 20 µM CNO for thirty minutes, TBS failed to induce LTP (Fig. 3A-B). The slopes of LTP time course following 1 µM and 20 µM CNO were significantly different (Linear regression followed by ANCOVA, *p* < 0.0001, n = 5-6/ dose). As 20 µM CNO administration for thirty minutes led to a decline in fEPSP slope, we re-analyzed the data by normalizing the fEPSP slope to ten minutes of pre-LTP data treated as baseline which showed a significant difference between slopes of LTP time course between 1 µM and 20 µM CNO treated slices (Fig. 3C, linear regression followed by ANCOVA, *p* < 0.027, n = 5-6/ dose). For further quantification, we binned the fEPSP slope data in five-minute intervals. Thirty minutes pre-treatment with 1 µM CNO led to a robust induction of LTP with the potentiated fEPSP slope in both 5-10 min and 25-30 min post-LTP time bins as compared to the pre-LTP time bin (Fig. 3D, post-hoc Bonferroni multiple comparisons following one-way ANOVA, *p* < 0.05, n = 5). This analysis also revealed a small, but significant increase in fEPSP slopes in 5-10 min and 25-30 min post-LTP time bins as compared to the pre-LTP time bin following administration of 20 µM CNO (Fig. 3E, post-hoc Bonferroni multiple comparisons following one-way ANOVA, *p* < 0.05, n = 6).

**Fig. 3.**
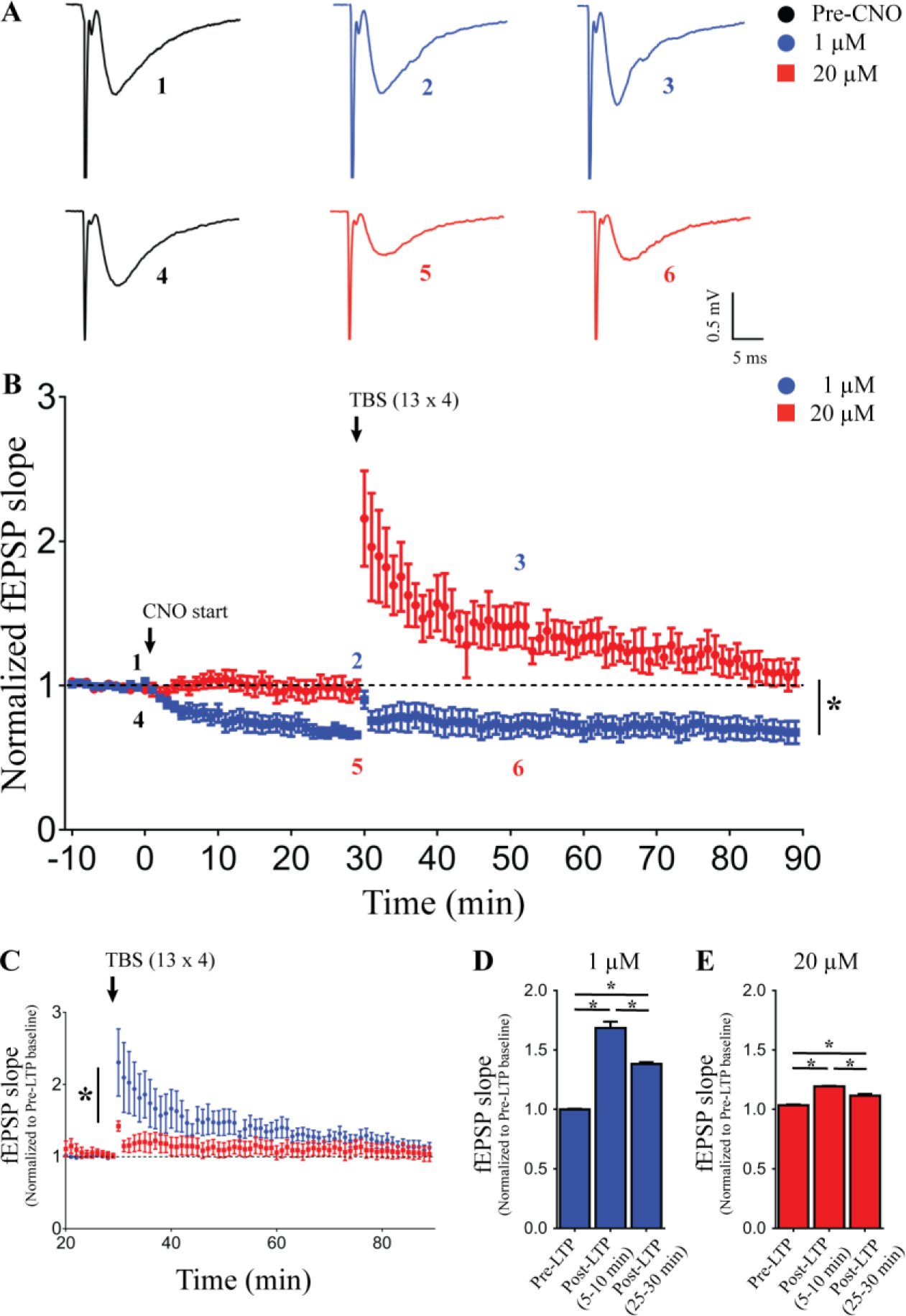
Dose-dependent effects of acute chemogenetic hM3Dq activation on theta burst stimulation induced LTP in hippocampal slices. (A) Shown are representative fESPS traces before, thirty minutes after CNO treatment and twenty minutes post LTP induction. Slices with thirty minutes of 1 µM CNO bath application show robust induction in LTP, with failure to induce LTP at 20 µM CNO treatment (B). fEPSP slope normalized to pre-LTP baseline shows robust potentiation with bath application of 1 µM CNO, but not at 20 µM CNO (C). The average fEPSP slope in five-minute time bins is significantly increased in 5-10 min and 25-30 min bin as compared to pre-LTP with 1 µM CNO bath application (D). The average fEPSP slope in five-minute time bins wass significantly increased in 5-10 min and 25-30 min bin as compared to pre-LTP with 20 µM CNO bath application (E). Results are expressed as the mean ± S.E.M. \**p* < 0.05 as compared between 1 µM and 20 µM CNO treatment (linear regression followed by ANCOVA) for time course analysis; \**p* < 0.05 as compared between pre-CNO, 1 µM and 20 µM CNO treatment (One-way ANOVA followed by post-hoc Bonferroni multiple comparison) for others.

These results demonstrate dose-dependent effects of CNO-mediated chemogenetic activation of CamKIIα-positive excitatory neurons on hippocampal plasticity, with slices pre-treated with low dose of CNO showing robust LTP induction, whereas TBS failed to induce LTP in slices exposed to a high dose of CNO.

### Acute CNO mediated hM3Dq DREADD activation of hippocampal excitatory neurons does not alter intrinsic membrane properties

We next performed whole cell patch-clamp on CA1 pyramidal neurons and investigated the effects of CNO administration on intrinsic membrane properties for a duration of thirty minutes. First, we generated a single action potential by injecting a step current of 2 nA for 2 ms which was used to confirm cell identity. We did not find any significant change in action potential threshold following thirty minutes of bath application of both 1 µM and 20 µM CNO (Table 1, n = 24 cells (pre-CNO), 6 cells (1 µM), 11 cells (20 µM)). As injection of a depolarizing pulse leads to firing of action potentials making a measurement of other membrane properties difficult, we next injected a hyperpolarizing step current of −100 pA for 500 ms and recorded the membrane voltage. We did not find any significant change in resting membrane potential (RMP), input resistance (R_N_), sag voltage and accommodation index following thirty minutes of CNO treatment (both 1 and 20 µM) as compared to the pre-CNO baseline (Table 1, n = 24 cells (pre-CNO), 6 cells (1 µM), 11 cells (20 µM)).

**Table. 1.** Acute CNO mediated hM3Dq DREADD activation of hippocampal excitatory neurons does not alter intrinsic membrane properties

| | RMP<br>(mV) | Input resistance<br>(MΩ) | AP threshold<br>(mV) | $\tau$ (ms) | Sag (mV) | Sag (%) | Accommodation<br>index |
| --- | --- | --- | --- | --- | --- | --- | --- |
| Pre-CNO | -61.31 ± 0.62 | 194.9 ± 10.95 | -53.61 ± 0.59 | 26.15 ± 1.57 | -5.67 ± 0.61 | 6.39 ± 0.61 | 0.45 ± 0.04 |
| 30 min<br>Post-CNO<br>(1 μM) | -59.24 ± 1.52 | 173.7 ± 17.47 | -56.04 ± 2.11 | 23.11 ± 1.66 | -4.84 ± 0.85 | 5.82 ± 0.84 | 0.36 ± 0.09 |
| 30 min<br>Post-CNO<br>(20 μM) | -62.30 ± 1.31 | 201.8 ± 13.21 | -53.39 ± 1.22 | 27.05 ± 3.03 | -4.71 ± 0.72 | 5.30 ± 0.74 | 0.47 ± 0.08 |

RMP: Resting membrane potential, AP threshold: Action potential threshold, $\tau$ : Membrane time constant

Taken together, these data show chemogenetic activation of CamKIIα-positive excitatory neurons has no significant influence on intrinsic membrane properties of CA1 pyramidal neurons following thirty minutes of either low or high dose CNO treatment.

### Bidirectional dose-dependent modulation of excitability following thirty minutes of hM3Dq chemogenetic activation of hippocampal excitatory pyramidal neurons

In order to understand possible dose-dependent effects of CNO mediated activation of hM3Dq DREADD in CamKIIα-positive excitatory neurons on intrinsic excitability, we injected increasing step currents and measured the number of action potentials (Fig. 4A). We found a significant increase in the number of action potentials produced in cells with thirty minutes of low dose (1 µM) CNO bath application (Fig. 4B-C, linear regression followed by ANCOVA, *p* = 0.04 (elevation), n = 24 cells (pre-CNO), 6 cells (1 µM), 11 cells (20 µM)). Further, there was a decrease in intrinsic excitability with thirty minutes of pre-treatment with high dose (20 µM) of CNO with a significant decline in number of action potentials generated by increasing amount of step current (Fig. 4B-C, linear regression followed by ANCOVA, *p* = 0.044 (slope), n = 24 cells (pre-CNO), 6 cells (1 µM), 11 cells (20 µM)). In order to understand the effects of chemogenetic activation on spontaneous activity of CA1 neurons, we measured spontaneous post-synaptic currents (sPSC) before and thirty minutes after activation CamKIIα-positive excitatory neurons using the low and high dose of CNO. We saw a significant difference in sPSC amplitude with thirty minutes of low dose (1 µM) CNO treatment with decrease in cumulative probability at lower amplitudes (< 35 pA) and increased cumulative probability at higher amplitudes (Fig. 4G, Kolmogorov-Smirnov two-sample comparison, *p* < 0.001, n = 13 cells (pre-CNO), 7 cells (1 µM)). However, we did not observe any significant change in sPSC inter-event interval (Fig. 4H, n = 13 cells (pre-CNO), 7 cells (1 µM)) following bath application of 1 µM CNO. Following thirty minutes treatment with high dose (20 µM) of CNO, we saw a significant decrease in both sPSC amplitude (Fig. 4I, Kolmogorov-Smirnov two-sample comparison, *p* < 0.001, n = 13 cells (pre-CNO), 7 cells (20 µM)) and inter-event interval (Fig. 4J, Kolmogorov-Smirnov two-sample comparison, *p* < 0.001, n = 13 cells (pre-CNO), 7 cells (20 µM)).

**Fig. 4.**
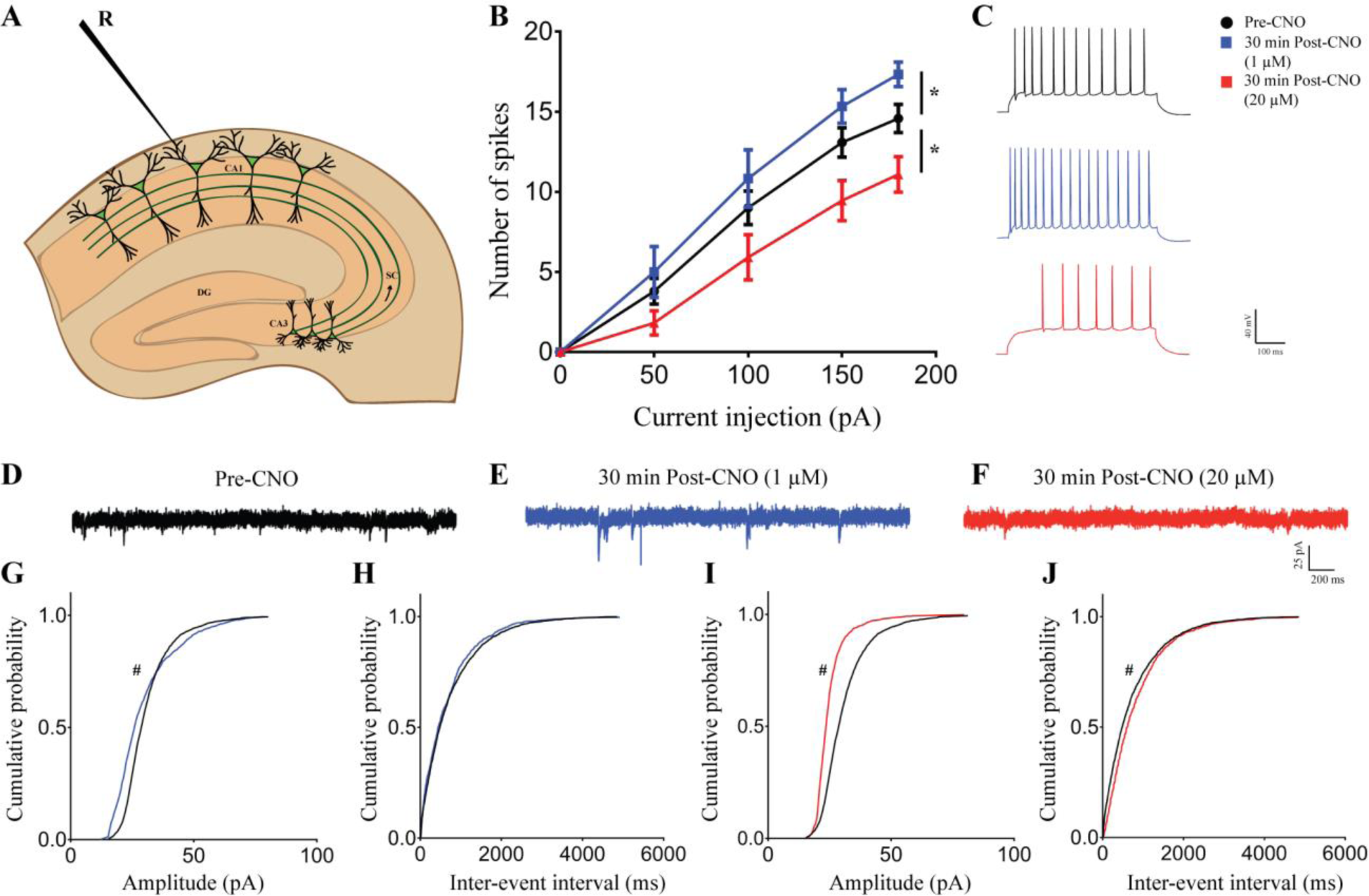
Bidirectional dose-dependent modulation of excitability following thirty minutes of hM3Dq chemogenetic activation of hippocampal excitatory pyramidal neurons. (A) Shown is a schematic depicting whole cell patch-clamp recording from somata of CA1 pyramidal cells. (B) The number of spikes generated with increasing amount of current injection is enhanced with bath application of 1 µM CNO but reduced with 20 µM CNO. (C) Shown is a representative trace of action potentials from CA1 neurons before CNO treatment, after thirty minutes bath application of 1 µM CNO, and 20 µM CNO. Results are expressed as the mean ± S.E.M. \**p* < 0.05 as compared between 1 µM and 20 µM CNO treatment (linear regression followed by ANCOVA). Shown are representative sPSC traces before (D) and thirty minutes after 1 µM (E) or 20 µM CNO treatment (F). Bath application of 1 µM CNO for thirty minutes resulted in significantly altered sPSC amplitude with decreased cumulative probability at lower amplitude and increased cumulative probability at higher amplitude (>35 pA) (G), whereas no significant effect was observed on inter-event interval (H). Bath application of 20 µM CNO for thirty minutes significantly decreased sPSC amplitude (I) and sPSC inter-event interval (J). Results are expressed as cumulative probabilities. # *p* < 0.001 as compared to pre-CNO treated group (Kolmogorov-Smirnov two-sample comparison). R – recording electrode.

This data suggests that intrinsic excitability of CA1 pyramidal neurons is bidirectionally regulated following CNO-mediated activation of CamKIIα-positive excitatory neurons, with low dose CNO (1 µM) increasing and high dose (20 µM) of CNO treatment decreasing excitability respectively. Further, the above data shows dose-dependent bidirectional regulation of spontaneous post synaptic current amplitude and frequency following CNO mediated activation CamKIIα-positive excitatory neurons, with low dose CNO increasing and high dose CNO mediated chemogenetic activation associated with decreased spontaneous activity.

### Dose-dependent effects of acute chemogenetic hM3Dq activation on AMPAR and NMDAR-mediated currents

We next sought to understand the possible cellular mechanism responsible for the decline in fEPSP and decreased synaptically-driven excitability (Fig. 2) of CA1 pyramidal cells at the higher dose of CNO treatment. As chemogenetic activation of CamKIIα-positive excitatory neurons would act by releasing glutamate and further action on glutamatergic receptors, we performed time course measurements of evoked AMPAR and NMDAR-mediated currents from CA1 pyramidal cells for thirty minutes following administration of either low (1 µM) or high dose (20 µM) of CNO (Fig. 5A). AMPAR-mediated current was calculated as the peak amplitude of evoked current when the cell was voltage clamped at −70 mV, and NMDAR-mediated currents were obtained as the average current 80-100 ms following the time of peak response when the cell was voltage clamped at +40 mV (Fig. 5B). AMPAR-mediated evoked current was reduced following both 1 and 20 µM CNO administration, but the extent of decline was lower in the 20 µM CNO administered neurons as compared to 1 µM CNO administered neurons (Fig. 5C,E linear regression followed by ANCOVA, *p* < 0.0001, n = 6 cells (1 µM), 5 cells (20 µM)). Similarly, we saw a reduction in NMDAR-mediated currents with both 1 and 20 µM CNO administration, and the magnitude of decline was higher in the 20 µM CNO administered neurons as compared to 1 µM CNO administered neurons (Fig. 5D, F, linear regression followed by ANCOVA, *p* < 0.0001, n = 5 cells (1 µM), 6 cells (20 µM)). To get a neuronal population activity, we compared average currents in five-minute bins before CNO treatment and thirty minutes following low and high dose CNO treatment. We found a significant decline in both AMPAR-mediated current (Fig. 5G, one-way ANOVA followed by Bonferroni multiple comparison, *p* < 0.05, n= 11 cells (pre-CNO), n = 6 cells (1 µM), 5 cells (20 µM)) and NMDAR-mediated current (Fig. 5H, one-way ANOVA followed by Bonferroni multiple comparison, *p* < 0.05, n= 11 cells (pre-CNO), n = 5 cells (1 µM), 6 cells (20 µM)) at both 1 and 20 µM CNO as compared to pre-CNO currents. Further, post-hoc Bonferroni multiple comparisons revealed that administration of 20 µM CNO significantly lowered both AMPAR-mediated current (One-way ANOVA followed by Bonferroni multiple comparison, *p* < 0.05) and NMDAR-mediated current (One-way ANOVA followed by Bonferroni multiple comparison, *p* < 0.05) as compared to 1 µM CNO treatment. Furthermore, the AMPAR/ NMDAR ratio was significantly lowered with thirty minutes of high dose (20 µM) CNO administration (Fig. 5I, one-way ANOVA followed by Bonferroni multiple comparison, *p* < 0.05, n = 8 cells (pre-CNO), 8 cells (1 µM), 7 cells (20 µM)). The time constant for NMDAR-mediated current decay was significantly lower following thirty minutes treatment of both 1 and 20 µM CNO (Fig. 5J, one-way ANOVA followed by Bonferroni multiple comparison, *p* < 0.05, n = 9 cells (pre-CNO), 5 cells (1 µM), 6 cells (20 µM)).

**Fig. 5.**
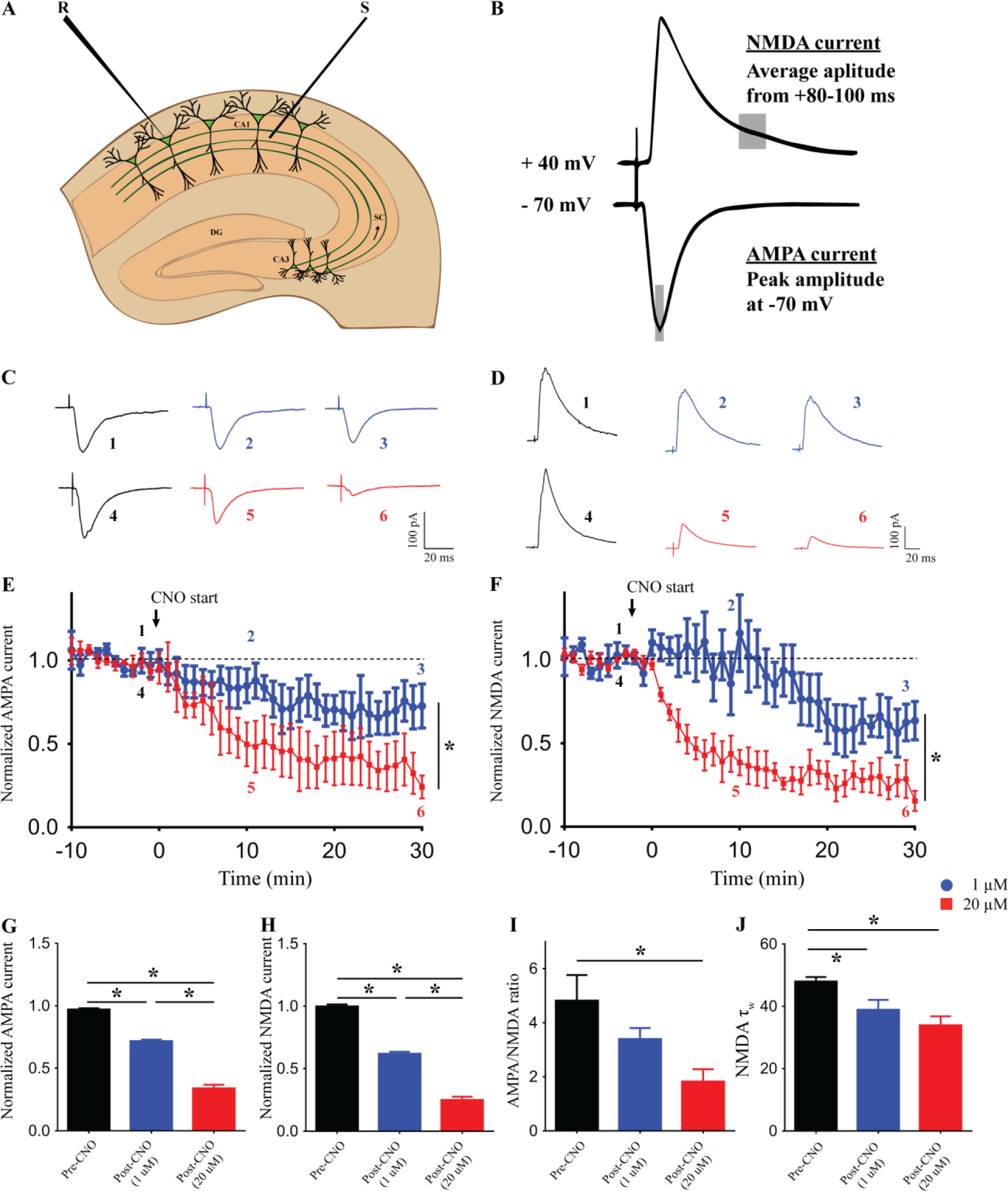
Dose-dependent effects of acute chemogenetic hM3Dq activation on AMPAR and NMDAR-mediated currents. (A) Shown is a schematic depicting whole cell patch-clamp recording from somata of CA1 pyramidal cells with stimulating electrodes places on Schaffer collateral input pathway. (B) Shown is a schematic demonstrating the measurement of AMPAR and NMDAR-mediated currents respectively. (C) Shown are representative AMPAR-mediated evoked post-synaptic current traces before, after 10 min and 30 min of CNO bath application. Voltage clamped at −70 mV. (D) Shown are representative NMDAR-mediated evoked post-synaptic current traces before, after 10 min and 30 min of CNO bath application. Voltage clamped at + 40 mV. Bath application of both 1 and 20 µM CNO results in reduced AMPAR-medaited currents, with significantly lower currents at 20 µM (E, G). Bath application of both 1 and 20 µM CNO results in reduced NMDAR-mediated currents, with significantly lower currents at 20 µM (F, H). AMPAR/ NMDAR ratio is significantly reduced with bath application of 20 µM CNO (I). The time constant of NMDAR-mediated current decay (τ) is significantly decreased following thirty minutes of both 1 and 20 µM CNO administration (J). Results are expressed as the mean ± S.E.M. \**p* < 0.05 as compared between 1 µM and 20 µM CNO treatment (linear regression followed by ANCOVA) for time course analysis; \**p* < 0.05 as compared between pre-CNO, 1 µM and 20 µM CNO treatment (One-way ANOVA followed by post-hoc Bonferroni multiple comparison) for others. S – stimulating electrode, R – recording electrode.

Taken together, these results indicate downregulation of the ionotropic glutamate receptor-mediated currents following chemogenetic activation CamKIIα-positive excitatory neurons by CNO in a dose-dependent manner.

### Acute administration of CNO results in a dose-dependent increase in intracellular calcium levels in primary hippocampal neurons

Upon CNO administration the Gq-coupled hM3Dq mobilizes intracellular calcium (Armbruster et al., 2007; Roth, 2016). As calcium and it’s downstream signaling are key regulators of neuronal physiology, we next sought to examine whether different doses of CNO lead to differential intracellular calcium dynamics using a ratiometric calcium-sensitive dye Indo1. Using two-photon calcium imaging, we measured intracellular calcium levels in primary hippocampal neurons following administration of either 1 or 20 µM of CNO. We observed a significant increase in calcium levels as noted from an increase in fluorescence intensity at both 1 µM (Fig.6A-B, linear regression, *p* < 0.0001) and 20 µM (Fig.6A-B, linear regression, *p* = 0.0002). Further, we also noted that administration of 20 µM CNO resulted in a significantly faster increase in intracellular calcium levels in the first ten minutes as compared to 1 µM CNO (Fig.6B, linear regression followed by ANCOVA, *p* = 0.012, n = 5 cells/ group, from 3 animals). Post ten minutes, the intracellular calcium reaches saturation level in cultured hippocampal neurons following acute CNO administration and was not significantly different between 1 µM and 20 µM of CNO.

**Fig. 6.**
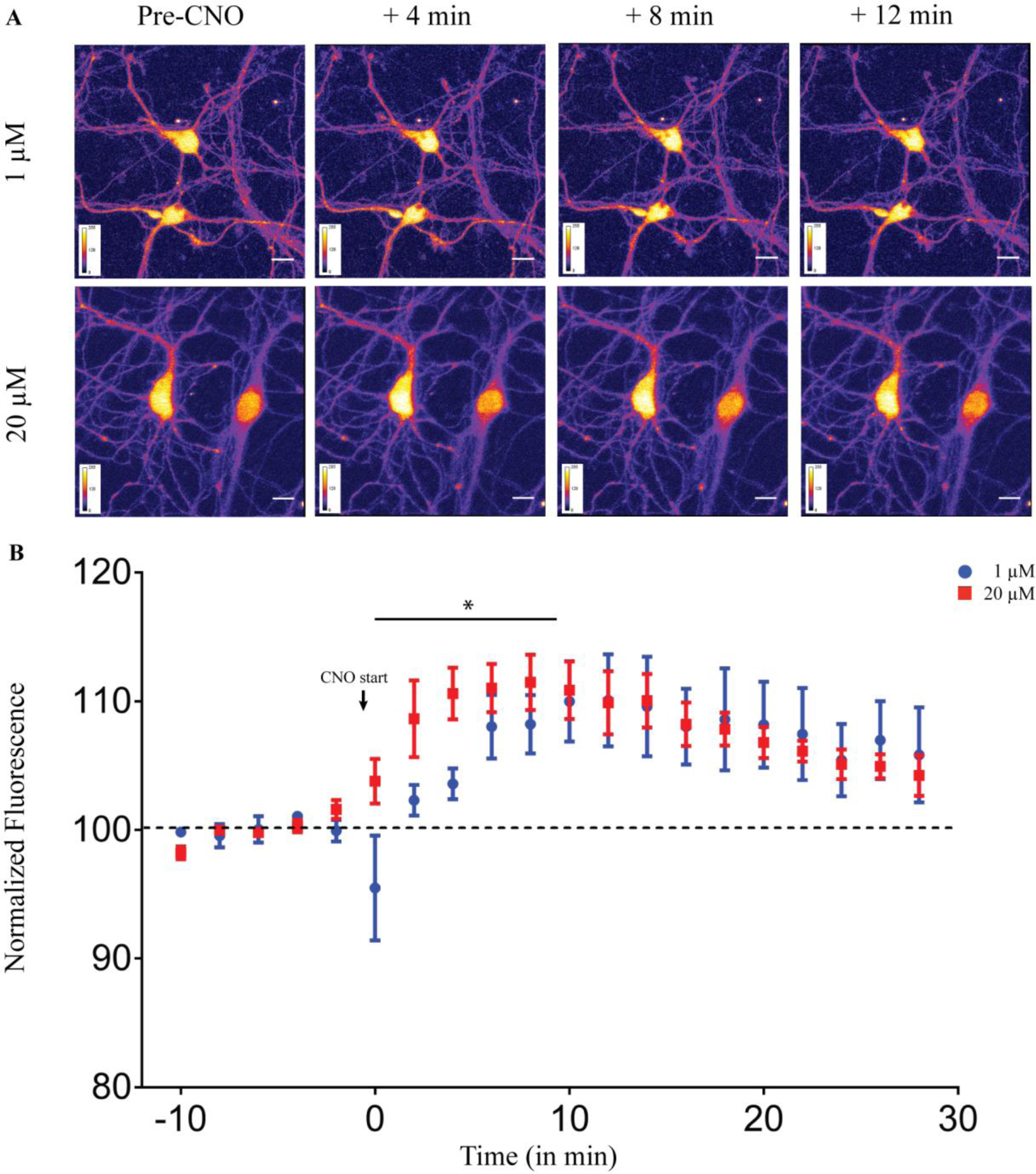
Acute administration of CNO results in a dose-dependent increase in intracellular calcium levels in primary hippocampal neurons. (A) Shown are representative images of primary hippocampal neurons before CNO-treatment and four, eight, and twelve minutes following 1 µM (top) or 20 µM (bottom) CNO treatment. (B) Application of both 1 µM and 20 µM CNO significantly increased intracellular calcium levels as observed by increased fluorescence intensity. Treatment with 20 µM CNO resulted in a significantly faster increase in intracellular calcium levels as compared to 1 µM CNO. Results are expressed as the mean ± S.E.M. \**p* < 0.05 as compared between 1 µM and 20 µM CNO treatment (linear regression followed by ANCOVA).

Taken together, our data show a dose-dependent increase in calcium levels following acute CNO administration in cultured hippocampal neurons with 20 µM CNO resulting in a significantly faster elevation in intracellular calcium levels than 1 µM CNO administration.

## Discussion

Our results provide novel insights into the dose-dependent regulation of hippocampal neurotransmission and plasticity by CNO-mediated activation of the hM3Dq DREADD in CamKIIα-positive excitatory neurons. We observed a significant decline in fEPSP slope in a dose-dependent manner with no effect at the low dose (1 µM) and the maximum decline noted at the high dose (20 µM) of CNO. Further, dose-dependent effects on hippocampal plasticity were indicated by a robust induction of TBS-evoked LTP in the presence of a low dose of CNO, which was lost at the high CNO dose. In keeping with this observation, the low dose of CNO potentiated PPR and increased excitability, accompanied by an intriguing regulation of the sPSC with enhanced probability for low amplitude events following CNO bath application. In contrast, we observed a significant decline in excitability and sPSC at the Schaffer collateral synapses, thirty minutes following bath application of a high dose of CNO. Taken together, our findings provide novel evidence for the complex nature of modulation of hippocampal neurotransmission following hM3Dq DREADD activation by different doses of the ligand, CNO.

The hM3Dq DREADD is a chemogenetic tool widely used to activate neurons in a circuit and cell type-specific manner that provides critical spatiotemporal control of neurocircuitry, thus facilitating the establishment of causal links between specific neuronal pathways and behavior (Roth, 2016). Application of the DREADD ligand CNO activates the canonical Gq-signaling pathway, increasing accumulation of inositol monophosphate (IP1) and enhancing intracellular Ca^2+^ levels (Alexander et al., 2009; Armbruster et al., 2007), and is thought to thus increase neuronal firing. Several studies have demonstrated increased firing rate and membrane depolarization in hippocampal CA1 pyramidal neurons (Alexander et al., 2009), raphe serotonergic neurons (Urban et al., 2016), and arcuate nucleus AgRP neurons (Atasoy et al., 2012; Krashes et al., 2011) in response to CNO-mediated hM3Dq activation. However, these studies employ a wide range of CNO doses ranging from 0.5 to 200 µM, and have performed recordings predominantly for short time scales of a few minutes following CNO administration (Hurni et al., 2017; Mahler et al., 2014). Given that most behavioral paradigms using acute hM3Dq DREADD activation operate across timescales of thirty minutes to several hours using diverse CNO dose ranges, it is critical to gain insight into the impact of CNO-based hM3Dq DREADD activation on neurotransmission, incorporating both different doses and longer time durations. Here, we show that CNO-mediated hM3Dq DREADD activation exerts complex, dose-dependent effects on hippocampal neurotransmission, including fEPSP slope, neuronal excitability, spontaneous and evoked activity, as well as plasticity.

Our results are in agreement with prior studies examining the influence of endogenous Gq-coupled GPCR activation on hippocampal neurotransmission. At the low dose of CNO (1 μM) mediated DREADD activation, our results agree with the increased excitability and potentiation of LTP phenotype noted with the M1 muscarinic agonist, carbachol that activates Gq signaling (Bröcher et al., 1992; Burgard and Sarvey, 1990; Natsume and Kometani, 1997). Prior evidence indicates that the mGluR1/5 agonist (R,S)-3,5-dihydroxyphenylglycine (DHPG)-dependent LTP is associated with increased excitability of CA1 pyramidal neurons and requires activation of protein kinase C, that lies downstream to Gq-signaling (Brager and Johnston, 2007). In addition, stimulation of the Gq-coupled muscarinic acetylcholine receptor 1 and 5 (mAChR; M1/5) leads to facilitation of LTP and synaptic transmission in both the hippocampus and visual cortex (Bröcher et al., 1992; Burgard and Sarvey, 1990; Natsume and Kometani, 1997). Our results with the low dose of CNO (1 µM) recapitulate several components of these phenomena associated with activation of either the M1/5 or group I mGluR in the hippocampus. In contrast, bath administration of both mGluR1/5 and mAChR; M1/5 agonists (Caruana et al., 2011; Cobb and Davies, 2005; Gladding et al., 2009; Kumar, 2010; Palmer et al., 1997) are also known to evoke long-term depression (LTD) at the Schaffer collateral synapses, which is similar to the effects observed with the high dose of CNO (20 µM) used to evoke hM3Dq DREADD activation. Concurring with this data, bath application of broad-spectrum mGluR agonists (±)-1-aminocyclopentane-trans-1,3-dicarboxylic acid (ACPD) (Schoepp et al., 1999), mGluR1/5 agonist DHPG (Schoepp et al., 1999), and mGluR5 specific agonist (R,S)-2-chloro-5-hydroxyphenylglycine (CHPG) (Fitzjohn et al., 1999; Huber et al., 2000; Palmer et al., 1997), also induces robust LTD in several brain regions, including the Schaffer collaterals (Bellone et al., 2008; Gladding et al., 2009). It is difficult to directly compare the doses of CNO used in our study to evoke hM3Dq DREADD activation, to the agonists used in prior studies to stimulate the mGluR1/5 and mAChR. It is rather striking that hM3Dq DREADD activation at distinct doses exerts effects similar to those reported with mGluR1/5 and mAChR stimulation, that are known to evoke differential effects on hippocampal neurotransmission, as well as divergent effects on plasticity. Our results suggest that the engineered hM3Dq DREADD, like its endogenous Gq-coupled receptor counterparts, can exert complex regulation on hippocampal neurotransmission.

Our data indicate that potentially both post-synaptic and pre-synaptic mechanisms are triggered following bath application of CNO, and these responses likely differ at across doses of CNO. Several studies demonstrate the cross-talk between Gq-coupled signaling pathway and ionotropic glutamate receptors (iGluRs) by both Ca^2+^-dependent and Ca^2+^-independent pathways mediated via scaffolding proteins like Homer and Shank (Gladding et al., 2009; MacDonald et al., 2007; Mao et al., 2005). Interestingly, mGluR-LTD is associated with downregulation of surface AMPAR-mediated by p38 mitogen activated protein kinase (MAPK) pathway (Huang et al., 2004; Moult et al., 2008; Rush et al., 2002) or the microtubule-associated protein 1B (MAP1B) mediated pathway (Davidkova and Carroll, 2007), as well as internalization of NMDA receptors from the synapses (Snyder et al., 2001). Corroborating with these data, we observe downregulation in both AMPAR and NMDAR-mediated currents following both low and high doses of CNO application, which support the possibility of postsynaptic modulation by CNO-mediated hM3Dq activation.

Further, we see a potentiation of PPR and increased excitability following thirty minutes of CNO bath application only at the low dose. Activation of Gq-coupled receptors increases intracellular Ca^2+^ (Berridge, 1993; Billups et al., 2006; Larkum et al., 2003; Nakamura et al., 2000), it is possible bath application of different dosage CNO could lead to different concentrations of intracellular Ca^2+^ both at the pre-synapse and post-synapse. Increase in Ca^2+^ in the pre-synapse could be responsible for elevated sPSC frequency and increased excitability, which could be potentially mediated by enhanced neurotransmitter release. Further, it is well characterized that different levels and temporal kinetics of intracellular concentration of Ca^2+^ are capable of recruiting distinct signaling pathways (Augustine et al., 2003; Berridge, 1998; Thomas et al., 1996). With a body of literature showing coupling of intracellular Ca^2+^ to Gq-coupled GPCR activation (Berridge, 2009, 1993; Billups et al., 2006; Larkum et al., 2003; Nakamura et al., 2000), differential responses including iGluR currents and sPSC can be attributed to differential intracellular Ca^2+^ dynamics. Interestingly, we observe a faster increase in intracellular Ca^2+^ levels at a higher dose of CNO in cultured hippocampal neurons. Further careful investigation is required to tease out the exact role of downstream Ca^2+^-mediated signaling in mediating complex regulation of neuronal physiology by activation of the hM3Dq DREADD.

Our data differs from the results of López et al. demonstrating enhanced LTP following bath application of CNO (5 µM) in C57Bl/6 animals virally expressing hM3Dq in CamKIIα-positive neurons in the CA1 (López et al., 2016). The differences in results could be due to the fact that the expression of hM3Dq was driven using viral versus bigenic mouse lines, and was spatially restricted to just the CA1 versus the entire hippocampus, in addition to other differences in age of animals used. This further substantiates the argument to be careful when interpreting neuronal activation/ behavioral data using chemogenetic tools. To rule out off-target effects of CNO and its metabolites, we performed fEPSP time-course measurements in hippocampal slices derived from the background C57Bl/6J mice lacking the hM3Dq receptor. We did not observe any significant effect both at 1 and 20 µM of CNO administration for one hour. Although our results ruled out any off-target effects on fEPSP timecourse in the Schaffer collaterals, further control experiments would be useful to rule out non-specific effects on other physiological measures.

In conclusion, our data elucidates that treating hM3Dq DREADD activation simply as neuronal excitation, misses the nuances of regulation of various aspects of neurotransmission over time scales and across dose ranges. It is possible that different doses of CNO can have very different pharmacological effects including recruitment of differential downstream signaling pathways, differential internalization kinetics, and cross talk with other ionotropic and metabotropic receptors. The signaling pathways recruited by activation of GPCRs like hM3Dq are arguably distinct from those recruited during the use of optogenetic methods to activate ion channels such as ChR. Although chemogenetic tools are invaluable in the quest for neuronal circuit/cell-specific modulation of behavior, our results motivate future experiments to carefully address the physiological effects of hM3Dq DREADD activation as an important component during interpretation of behavioral data.

## Author contributions

SP, JCL, and VV designed the experiments. SP carried out slice physiology experiments. SS performed immunofluorescence staining to detect HA-tag. SS, and AB performed primary cortical neuron cultures. SP, MK and SM performed and analyzed ratiometric two-photon calcium imaging. SP and VV wrote the paper.

## Acknowledgments

We acknowledge Dr. Shital Suryavanshi and Dr. R.G. Prakash for their help and assistance with animal work.

