## Supplemental figures for "Chemogenetic activation of excitatory neurons alters hippocampal neurotransmission in a dose-dependent manner"

### Extended Figures

### Supplementary Fig. 1.

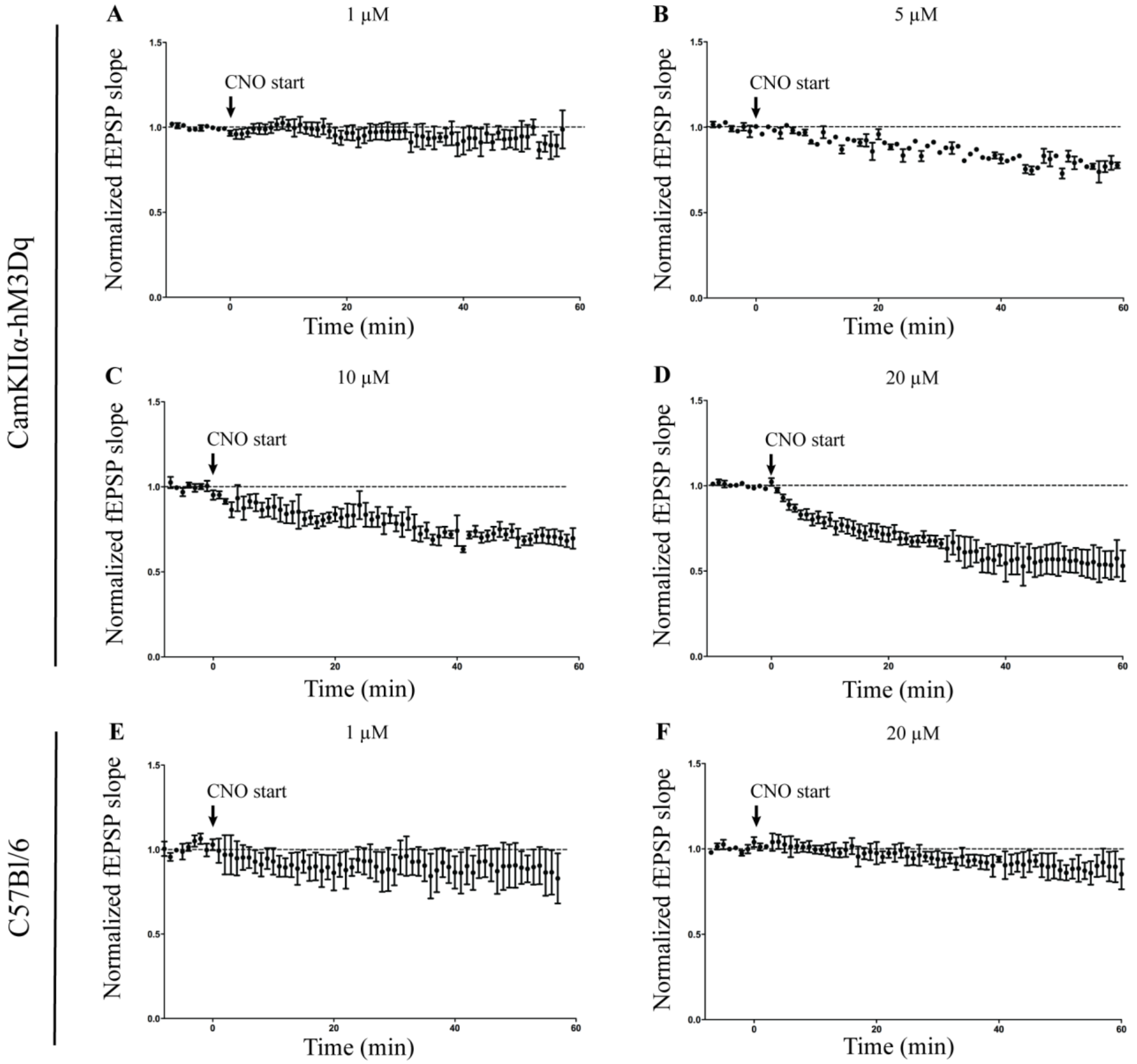


Bath application of hM3Dq DREADD agonist CNO had no effect on fEPSP at low dose (1 μM; A). CNO treatment induced long-term depression at a subsequently high dosage of 5 μM (B), 10 μM (C), and 20 μM (D). No effect on fEPSP was observed at 1 (E) and 20 μM (F) of CNO in slices taken from background C57Bl/ 6J animals. Results are expressed as the mean ± S.E.M.

**Supplementary Fig. 2.**

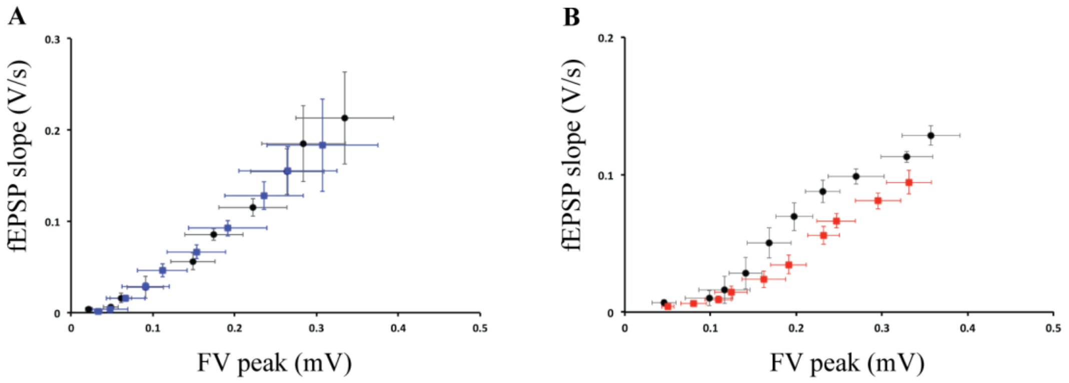


FV peak vs fEPSP slope at 1 µM (A) and 20 µM (B). Results are expressed as the mean ± S.E.M. *p < 0.05 as compared between 1 µM and 20 µM CNO treatment (linear regression followed by ANCOVA).
